# Sleep quality and quantity and their associations with age, salivary cortisol and abnormal behaviours: a preliminary field study in racing thoroughbreds

**DOI:** 10.64898/2026.01.31.702987

**Authors:** Noémie Hennes, Linda Greening, Sebastian McBride, Julie Lemarchand, Juliette Cognié, A Foury, Romane Phelipon, Hélène Bourguignon, Alice Ruet, Léa Lansade

## Abstract

Sleep plays a key role in both physical recovery and welfare. However, sleep patterns remain poorly documented in animals, particularly in athletic horses. This study aimed to provide a detailed description of sleep quantity and quality in training Thoroughbred racehorses and to investigate their relationships with age, abnormal behaviours, and cortisol. Thirteen Thoroughbreds (2–7 years old) were continuously monitored in their home environment over three consecutive days. An ethogram was used to quantify the two main phases of sleep: Non Rapid Eye Movement sleep (NREM) and Rapid Eye Movement sleep (REM), as well as sleep interruptions (from day 1 at 12:00 a.m. to day 3 at 12:00 a.m.). Sleep Quality Indices (SQI), defined as the quantity of sleep divided by the number of sleep interruptions (SI), were calculated. Behavioural observations of four indicators of poor welfare (alertness, stereotypies, inactivity, aggressiveness towards humans) were performed using scan sampling, and salivary cortisol was measured each morning. Linear models were used to assess the links between sleep quantity and quality, age, mean cortisol, and abnormal behaviours. Sleep quantity was significantly associated with age: positively for total NREM sleep (ANOVA: χ² = 5.26, p < 0.05) and, negatively for total REM sleep (ANOVA: χ² = 4.46, p <0.05) and total recumbency duration (ANOVA: χ² = 5.68, p < 0.05), suggesting an age-related shift favouring NREM over REM. Morning cortisol concentrations and the frequency of abnormal behaviours were significantly higher in horses with lower sleep quality (cortisol: Total SQI, ANOVA: F = 5.26, p < 0.05; Combined SQI, ANOVA: F = 5.40, p < 0.05; abnormal behaviours: Total SQI, ANOVA: F = 4.07, p = 0.074), pointing to a potential link between stress or altered welfare and poorer sleep quality. These findings suggest that, whilst the type and duration of equine sleep may be mainly affected byage, sleep quality is associated with both cortisol levels and the expression of abnormal behaviours, indicating that poor sleep quality may be linked to poor welfare in this population of horses. Thus, sleep appears to be closely linked with racehorse welfare, highlighting the need for further investigation into how it is influenced by factors such as husbandry, training load, recovery, and performance.

## Introduction

Sleep is a fundamental need for mammals. It consists of two main phases: Non-Rapid Eye Movement sleep (NREM) also known as slow-wave sleep; and Rapid Eye movement sleep (REM) also referred to as paradoxical sleep. NREM and REM phases alternate in cycles. The NREM phase is involved in different functions, such as muscle recovery and energy conservation while REM sleep plays an essential role in memory consolidation and cognitive processing [1]. Each species has their own sleep patterns which can vary depending on life stage, with young individuals commonly sleeping more than adults, especially in the REM phase [2,3]. Because of its essential role in recovery, sleep has been extensively studied in human athletes [4–7], who are considered to have greater sleep needs than sedentary individuals [7]. Appropriate sleep duration appears to support sports performance as well as overall well-being, whereas lack of sleep is associated with negative mood [8,9], higher risk of injuries [5,10] and poor athletic performance [9]. For instance, 48 hours of sleep deprivation after muscle damage induced by eccentric exercise resulted in higher cortisol levels than the same individuals under normal sleep conditions [6], and in swimmers, cortisol concentrations before major competition were positively associated with fatigue scores [11]. Moreover, sleep disturbances are observed in athletes experiencing non-functional overreaching or overtraining, along with other physiological and behavioural symptoms [12]. This reinforces the idea that sleep is essential for both well-being and performance in athletes, but also that sleep quantity and quality can be affected by the psychological and physiological consequences of sporting and athletic activity s[13,14].

Beyond its specific importance for athletic performance, sleep is also essential to overall human and non-human welfare. For example, in rodents, transitory sleep deprivation affects physiology as it enhances oxidative stress [15] and pain perception in rodents [16]. Less resting time during daytime or less lying time has been associated with higher expression of repetitive behaviours in dogs [17] and horses [18], suggesting that there is an association between sleep and abnormal behaviours in those two species. Housing conditions also influence sleep since restricted housing seems to negatively affect sleep quantity in rats [19], while sub-optimal conditions have been associated with shorter duration of recumbent sleep and longer duration of standing sleep in horses [20]. Additionally, horses tend to take longer time to solve a spatial memory task after 72 hours of REM sleep deprivation [21], suggesting that a lack of REM sleep could affect cognitive abilities in horses. Although many of these studies have focused on sleep duration, there is increasing consideration that sleep quality may be a better determinant and predictor of both welfare and physical/psychological performance [22]., In this respect, measuring sleep fragmentation, number of awakenings, or sleep efficiency may provide greater insights into actual sleep experience and how this relates to measures of wellbeing and performance. For instance, in humans, increased night-time awakenings has been associated with poor self-reported sleep quality, even when sleep duration appeared adequate [23]. In dogs, a higher number of awakenings has been linked to greater sensitivity to sad voices, considered a negative stimulus, during an emotion recognition test [24].

The aim of this study, therefore, was to investigate the relationship between sleep quantity/ quality and routinely used behavioural (e.g. inactivity, stereotypies, aggression) (Fureix and Meagher, 2015; Roberts et al., 2017; Fureix et al., 2010) and physiological (salivary cortisol) (König et al., 2017) markers of stress in the athletic horse (racing thoroughbreds). We also included the age of the horse within the analyses and hypothesized that total sleep, and particularly REM sleep, would vary with age, and that horses exhibiting more abnormal behaviours and higher morning cortisol levels would also show lower sleep quantity and potentially poorer sleep quality.

## Materials and methods

### Animals

Thirteen Thoroughbred racehorses (7 mares, 2 stallions and 4 geldings), aged between 2 and 7 years (mean ± SD: 3.46 ± 1.51), were included in the study. All horses had been in race training for at least six months and were housed at the same training centre in Normandy, France, under the care of two different trainers (7 horses for trainer A and 6 horses for trainer B). None of the horses had participated in a race during the four days prior to data collection.

Each horse was kept in an individual stable measuring 3.5 m × 4 m, with deep straw bedding (≥5cm), and was housed continuously in this stall, except during training sessions and routine management procedures. Each horse had *ad libitum* access to hay and water and were fed additional concentrate food in quantities adapted to their training needs, as determined by the trainer. Horses were exercised daily in sessions adjusted according to their training programme and fitness level, for approximately one hour per day between 06:15 and 12:00 a.m. A standardised veterinary health check was conducted on all horses on the first day of data collection, including assessments of intestinal sounds, heart rate, respiratory rate, mucous membrane colour, and lameness (SI, Table 1.). All individuals were assessed as clinically healthy.

**Table 1.** Overview of the measurement schedule over the three-day study period.

| Measured variable | Method | Timing | Duration |
| --- | --- | --- | --- |
| Salivary cortisol | Saliva sampling | Each morning (Days 1 to 3) | Once a day (3 days) |
| Sleep (quality, quantity) | Continuous video observation | Day 1 (12:00 am) to Day 3 (12:00 am) | 48h |
| Abnormal behaviours | Scan sampling (every 15 min) | Day 1 (12:00 am) to Day 3 (12:00 am) | 48h |

### Study schedule

The study was conducted over three consecutive days for each training yard, in autumn 2024. Salivary cortisol concentrations were collected each morning during the three-day period. Sleep quantity and quality, as well as abnormal behaviours, were assessed from video recordings obtained between Day 1 at 12:00 and Day 3 at 12:00 (48 h), without altering the horses’ usual routine (Table 1).

### Salivary cortisol

Salivary samples were collected daily by a trained experimenter, before exercise, in the early morning, between 06:15 and 08:15, using swabs (SARDTEDT cortisol Salivette®). Saliva was collected by gently moving the swab around the oral cavity, both above and below the tongue, for approximately 30 seconds to ensure it was sufficiently soaked. Samples were immediately frozen at −80 °C and stored until cortisol analysis.

Salivary cortisol concentrations were quantified from 50 µl saliva samples using a commercially available enzyme-linked immunosorbent assay (Cortisol Saliva ELISA; IBL International GmbH, Hamburg, Germany). The assay sensitivity was 0.05 ng/ml. Intra-assay and inter-assay coefficients of variation were 4.3% and 13.2%, respectively.

### Sleep assessment

Continuous video recordings were collected using two infrared wide-angle security cameras (Reolink E1 outdoor, 5MP PTZ) installed in each stall to cover it entirely. Recording started at 12:00 on day 1 and ended at 12:00 on day 3, allowing the capture of two complete nights and beginning/ending during a calm period of the day. This provided 48 hours of recording per horse, which is considered sufficient to assess sleep patterns in the absence of environmental changes [21]. Sleep behaviour was analysed using the behavioural ethogram developed by Greening and colleagues [22], which defines four sleep states: standing NREM, sternal NREM, sternal REM, and lateral REM sleep (Table1). Video analysis was performed continuously by two trained observers following a prior training phase to ensure consistency, using BORIS observer software [25]. Observers also reviewed recordings to identify and count sleep interruptions (SI) lasting more than 3 seconds and less than 3 minutes in each sleep phase, according to definitions by Greening et al. [20,22]. Based on these data, total sleep time (TST), total NREM sleep, total REM sleep, and total recumbency (REM and NREM recumbent sleep) durations were calculated over the 48-hour period.

Sleep quality was then assessed using five indices defined by Greening and colleagues [22]: Total Sleep Quality Index (SQI), Combined SQI, Weighted Combined SQI, NREM SQI and REM SQI as follows:

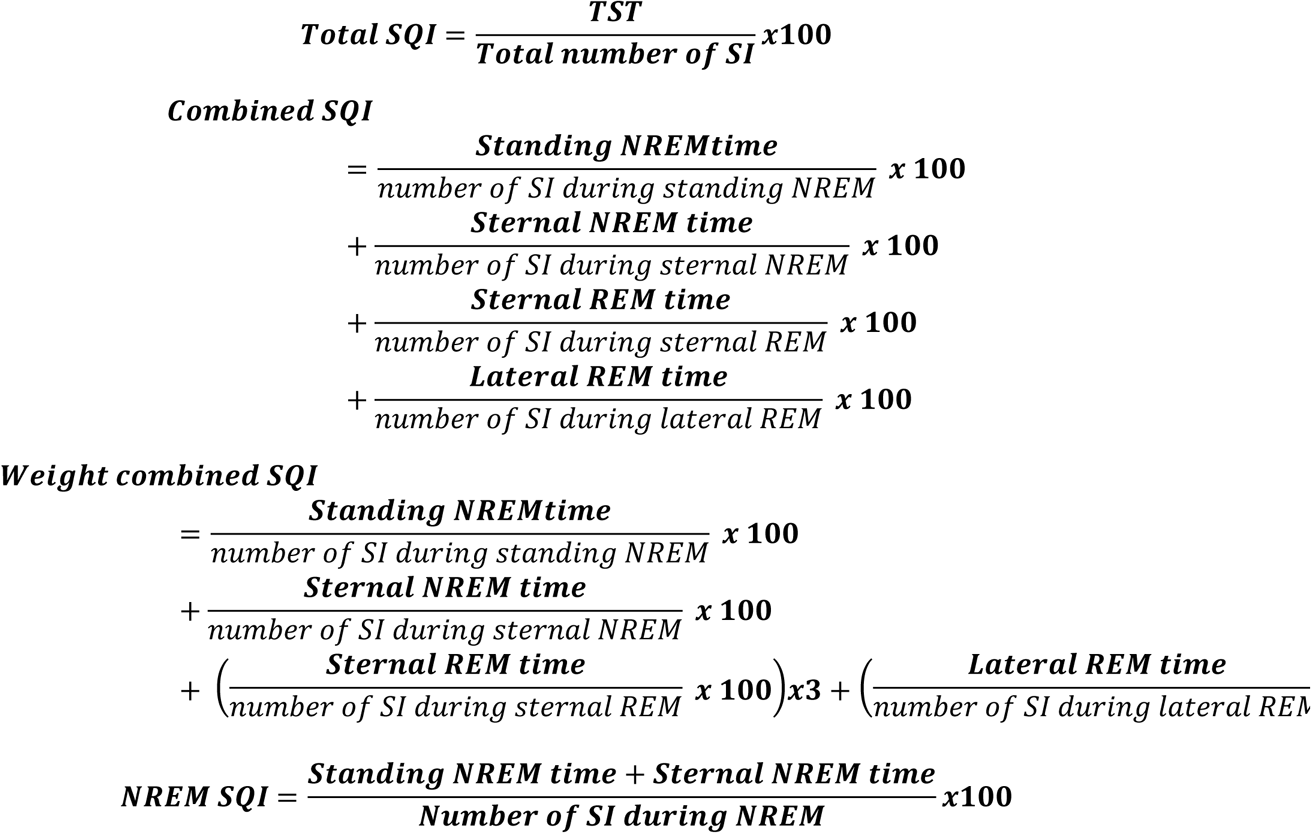

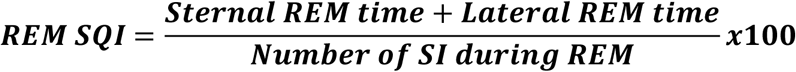

**Table 1.**
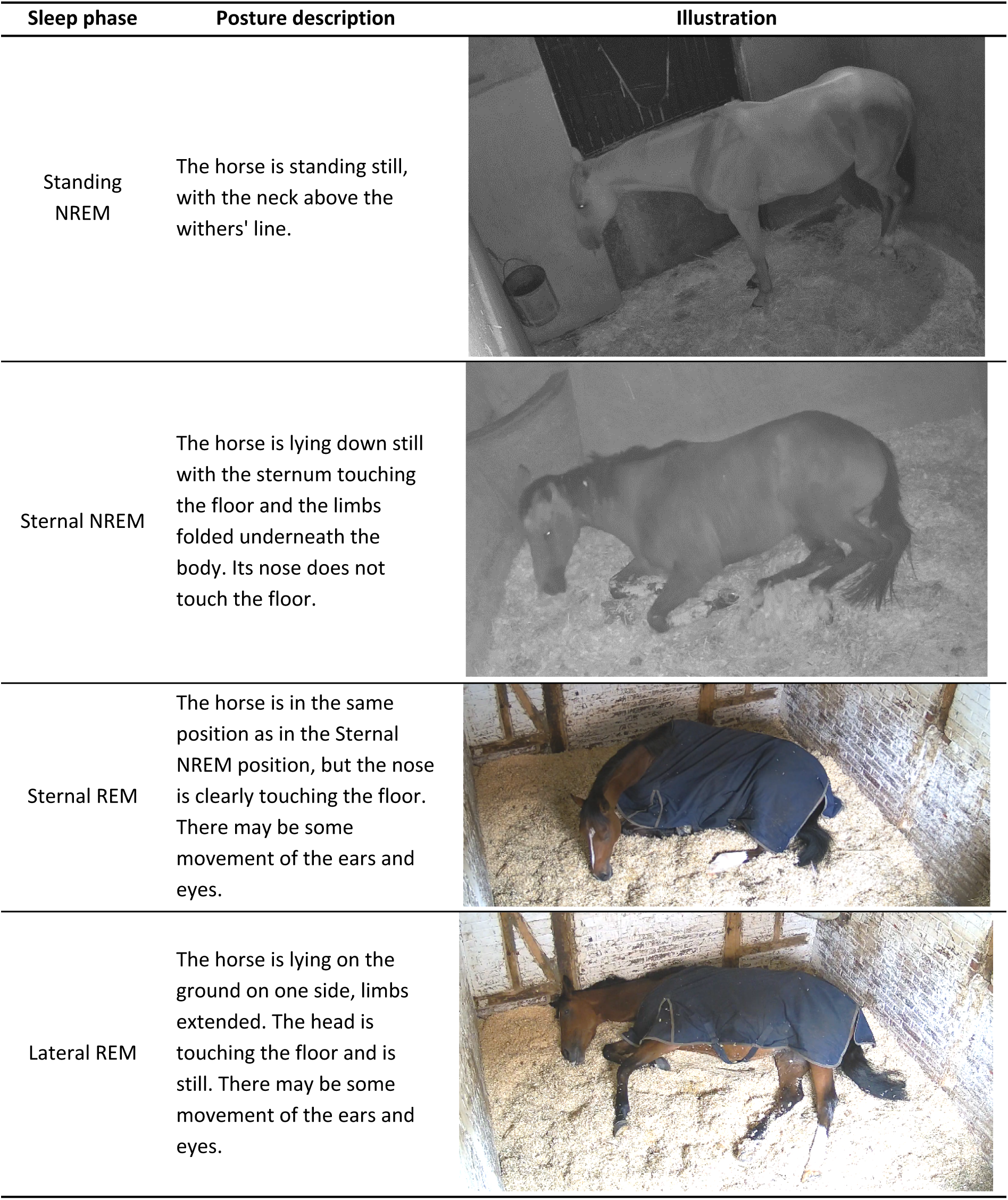
Ethogram of the different stages of sleep, based on that described by Greening and colleagues [20,22]. NREM = Non-Rapid Eye Movement, REM = Rapid Eye Movement.

| Sleep phase | Posture description | Illustration |
| --- | --- | --- |
| Standing NREM | The horse is standing still, with the neck above the withers' line. |  |
| Sternal NREM | The horse is lying down still with the sternum touching the floor and the limbs folded underneath the body. Its nose does not touch the floor. |  |
| Sternal REM | The horse is in the same position as in the Sternal NREM position, but the nose is clearly touching the floor. There may be some movement of the ears and eyes. |  |
| Lateral REM | The horse is lying on the ground on one side, limbs extended. The head is touching the floor and is still. There may be some movement of the ears and eyes. |  |

### Abnormal behaviours assessment

Abnormal behaviours monitored included inactivity, stereotypies, alert posture and aggression towards humans, as defined by other studies (Hennes et al., 2025; Phelipon et al., 2024; Ruet et al., 2022a) (Table 2) and considered as indicators of poor welfare [28–30]. Scan sampling was conducted every 15 minutes throughout the 48-hour observation period using video recordings. At each scan, the presence or absence of at least one of these behaviours was recorded. The number of observations per horse was 182.3 ± 4.1. Missing scans corresponded to periods when the horse was not in the stall, either due to training or for routine care. Then, the percentage of observations during which each horse displayed an abnormal behaviour was used for the following analyses. It was calculated as:

**Table 2.** Ethogram of four abnormal behaviours recorded using scan sampling, for each horse, according to Hennes et al. (2025) and Ruet et al. (2022).

| Abnormal behaviours | Description |
| --- | --- |
| Inactivity | The horse remains still on all four limbs, staring with eyes open and ears positioned either sideways or facing backward. The head and neck are carried at no more than 30° above the horizontal line through the withers, or below that horizontal line. During this posture, the horse is neither resting nor observing its environment. |
| Stereotypies | Stereotypic behaviours involving locomotion or oral activity, including repeated lip or tongue movements, weaving, repetitive head nodding or pacing. |
| Alert posture | The horse holds its neck elevated, with ears directed forward. This posture reflects heightened environmental awareness and a state of hypervigilance. |
| Aggressiveness towards humans | The horse displays aggressive behaviour, such as threats with flattened ears, or more direct actions like biting or kicking a human passing in front of the stall or entering the stall. |

% of abnormal behaviours = (Number of scans with any abnormal behaviours/ Total number of scans) x 100.

## Statistical analyses

All statistical analyses were conducted using RStudio software [31].

To investigate the effects of age, average morning cortisol levels and the expression of abnormal behaviours on TST, total REM sleep, total NREM sleep, total recumbency, and the five sleep quality indices, we fitted a linear model for each sleep-related variable. Fixed effects included age, the average morning cortisol level, calculated as the mean of the three morning samples, and the percentage of scans during which the horse exhibited an abnormal behaviour. Morning cortisol concentration (0.88 ± 0.44 ng/mL, range 0.23 to 2.35 ng/mL) did not vary significantly across the three days of the study (Friedman test: chi² = 1.08, df = 2, p = 0.58), supporting the use of their mean for the following analyses. Trainer was included as a random effect in all models, except for REM SQI, for which a model without a random effect was used, as no data transformation allowed model assumptions to be satisfactorily met when including a random effect.

Models were fitted using the lmer function from the lme4 package [32]. Model assumptions (normality and homoscedasticity of residuals) were assessed using simulated residual diagnostics from the DHARMa package [33]. When deviations from these assumptions were detected, appropriate transformations of the response variables (logarithmic or inverse transformations) were applied to improve model fit. The transformation applied to each response variable is reported in Table 5. Statistical significance was set at p ≤ 0.05.

An analysis of variance (ANOVA) was then performed on each model to determine which fixed factors significantly influenced the dependent variable. For mixed-effects models, the significance of fixed effects was assessed using Wald chi-square tests derived from Type II ANOVA.

## Results

### Descriptive results

#### Sleep pattern

The horses’ total sleep time (TST) averaged (±SD) 551.5 ± 110.9 minutes over the 48-hour video recording period, including 382.6 ± 101.2 minutes of NREM sleep and 168.9 ± 119.9 minutes of REM sleep (Fig.1, Tab. 3). Most sleep occurred in a standing position, followed by sternal NREM sleep. On average, lateral and sternal REM sleep were approximately equivalent in duration (Table 3). One gelding (4 years old) displayed exceptionally high values for both TST and REM sleep, with 469.2 minutes of REM sleep over 48 hours which represents nearly 2.7 times the group average. Conversely, one mare (4 years old) was never observed sleeping in lateral recumbency but did exhibit REM sleep while in sternal recumbency.

**Fig. 1:**
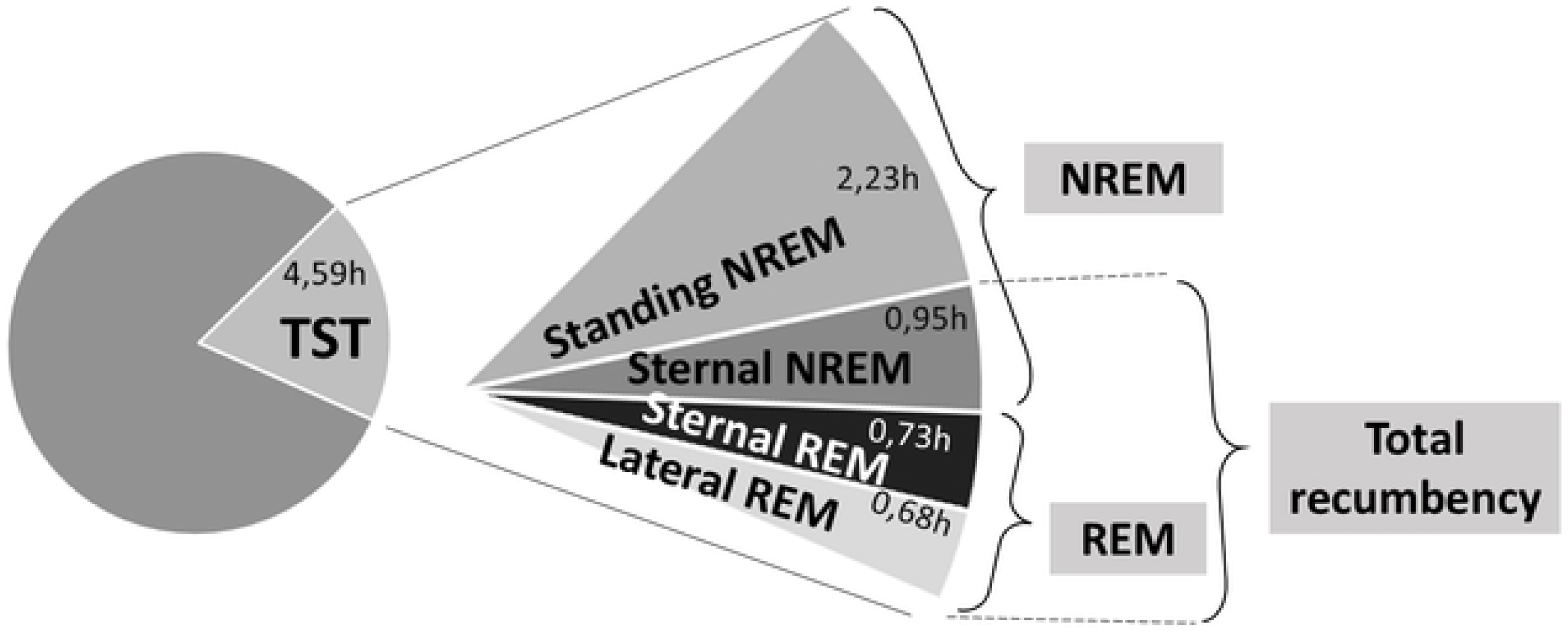
Graphical representation of the time spent sleeping in each position and sleep stage for the horses in this study. Sleep durations are expressed in hours per day and represent the average across all horses. *Data were recorded over a 48-hour period. TST = Total Sleep Time, NREM = Non-Rapid Eye Movement, REM = Rapid Eye Movement*.

**Table 3.** Description of the duration of each sleep phase and the number of sleep interruptions within each phase.

| Sleep phase | Duration of sleep (minutes) |  | Number of Sleep Interruptions (SI) |  |
| --- | --- | --- | --- | --- |
| | Mean±SD | Range | Mean±SD | Range |
| **Total Sleep Time** | $551.5 \pm 110.9$ | 386.5 – 781.0 | $76.2 \pm 17.3$ | 58 – 115 |
| **Standing NREM** | $268.6 \pm 103.1$ | 129.1 – 507.8 | $45.5 \pm 15.2$ | 15 – 79 |
| **Sternal NREM** | $113.9 \pm 73.3$ | 5.59 – 275.0 | $13.6 \pm 11.4$ | 0 – 37 |
| **Total NREM** | $382.6 \pm 101.2$ | 207.6 – 570.4 | $59.1 \pm 14.1$ | 37 – 84 |
| **Sternal REM** | $87.2 \pm 48.8$ | 27.1 – 215.3 | $13.8 \pm 10.8$ | 2 – 35 |
| **Lateral REM** | $81.8 \pm 82.4$ | 0.0 – 253.9 | $3.3 \pm 4.9$ | 0 – 16 |
| **Total REM** | $168.9 \pm 119.9$ | 68.7 – 469.2 | $17.2 \pm 15.1$ | 2 – 51 |
| **Total recumbency** | $282.9 \pm 142.6$ | 74.3 – 629.2 | $30.8 \pm 18.0$ | 9 – 64 |

#### Abnormal behaviours

All four types of abnormal behaviours were observed, although aggression towards humans was recorded only once in a single horse. The most commonly observed abnormal behaviour was stereotypy, with 7 out of 11 horses displaying such behaviours at least once. Specifically, four horses were observed performing repetitive headshaking, while three exhibited locomotor stereotypies, consisting of pacing in circles or in a figure-of-eight pattern. Five horses were observed at least once in an inactive posture, and four at least once in an alert posture. Overall, the mean total percentage of observations showing abnormal behaviours was 2.0 ± 2.5% (Table 4).

**Table. 4:**
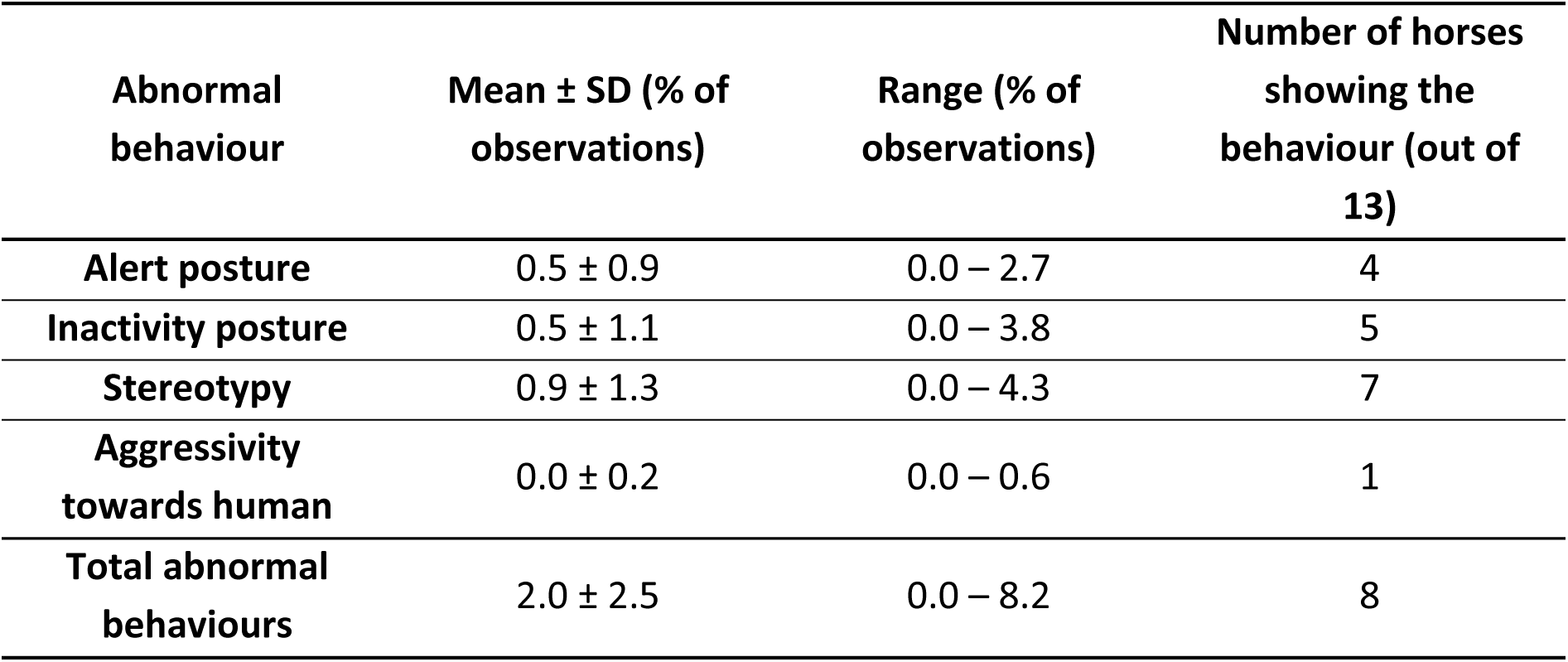
Percentages of observations and number of horses observed at least once expressing abnormal behaviours over the 48-hour recording period, using the scan sampling method with one observation every 15 minutes.

| Abnormal behaviour | Mean ± SD (% of observations) | Range (% of observations) | Number of horses showing the behaviour (out of 13) |
| --- | --- | --- | --- |
| Alert posture | $0.5 \pm 0.9$ | 0.0 – 2.7 | 4 |
| Inactivity posture | $0.5 \pm 1.1$ | 0.0 – 3.8 | 5 |
| Stereotypy | $0.9 \pm 1.3$ | 0.0 – 4.3 | 7 |
| Aggressivity towards human | $0.0 \pm 0.2$ | 0.0 – 0.6 | 1 |
| Total abnormal behaviours | $2.0 \pm 2.5$ | 0.0 – 8.2 | 8 |

**Table 5.**
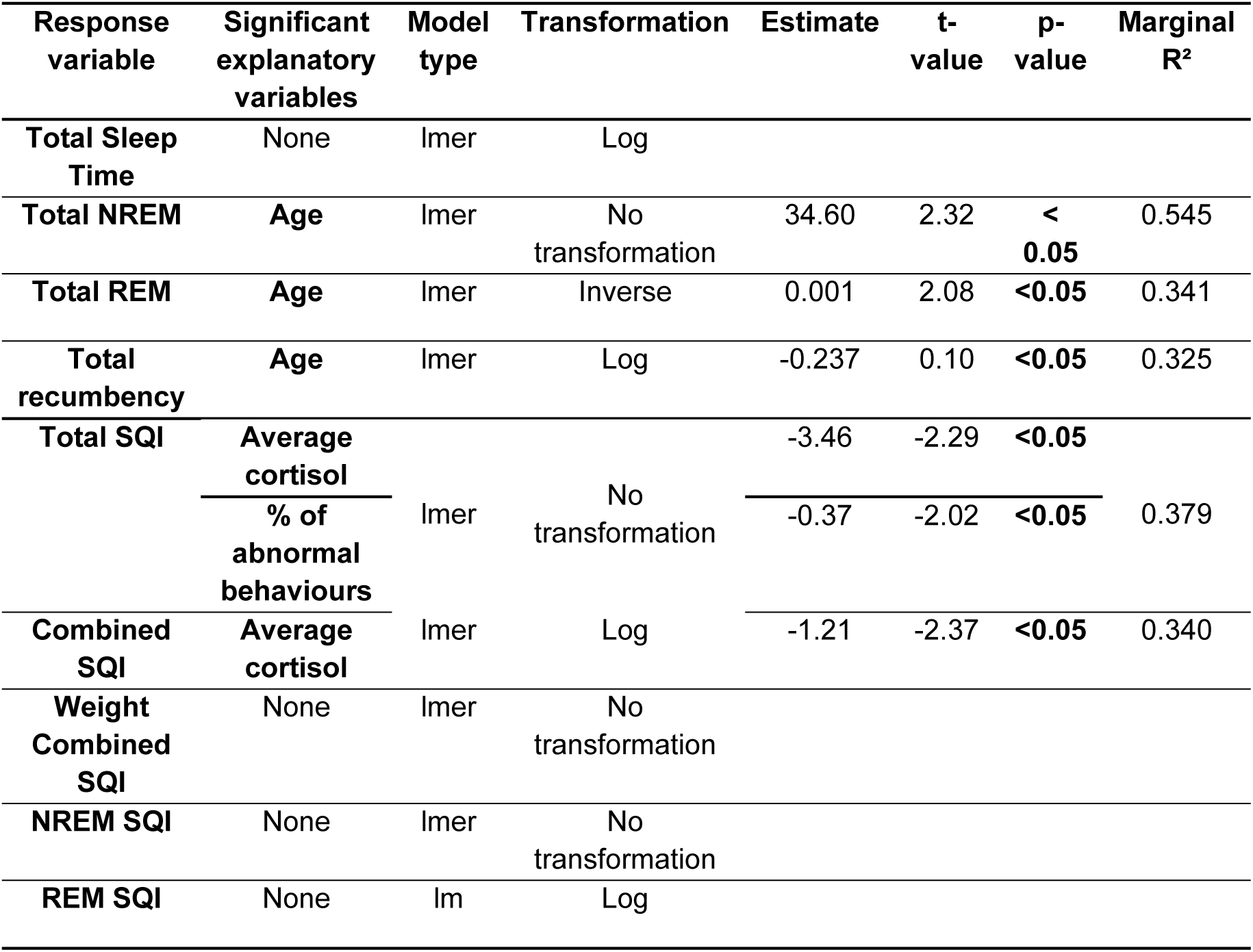
Summary of the results of the models on the different sleep quality and quantity variables with age, percentage of observations of abnormal behaviours, and average morning cortisol concentration as fixed factors. “lm”: linear models. NREM = Non Rapid Eye Movement, REM = Rapid Eye Movement.

#### Relationships between quantity of sleep, age, salivary cortisol, and abnormal behaviours

Total NREM sleep was positively associated with age (ANOVA: χ² = 5.4, p < 0.05; Fig. 2a), while total REM sleep and total recumbency were negatively associated with age (ANOVA: χ² = 4.4, p < 0.05; Fig. 2b and χ² = 5.7, p < 0.05; Fig. 2c). The models yielded marginal R² values of 54.5%, 34.1%, and 32.5%, respectively. Neither abnormal behaviours nor cortisol concentrations showed significant effects in the models.

**Fig. 2:**
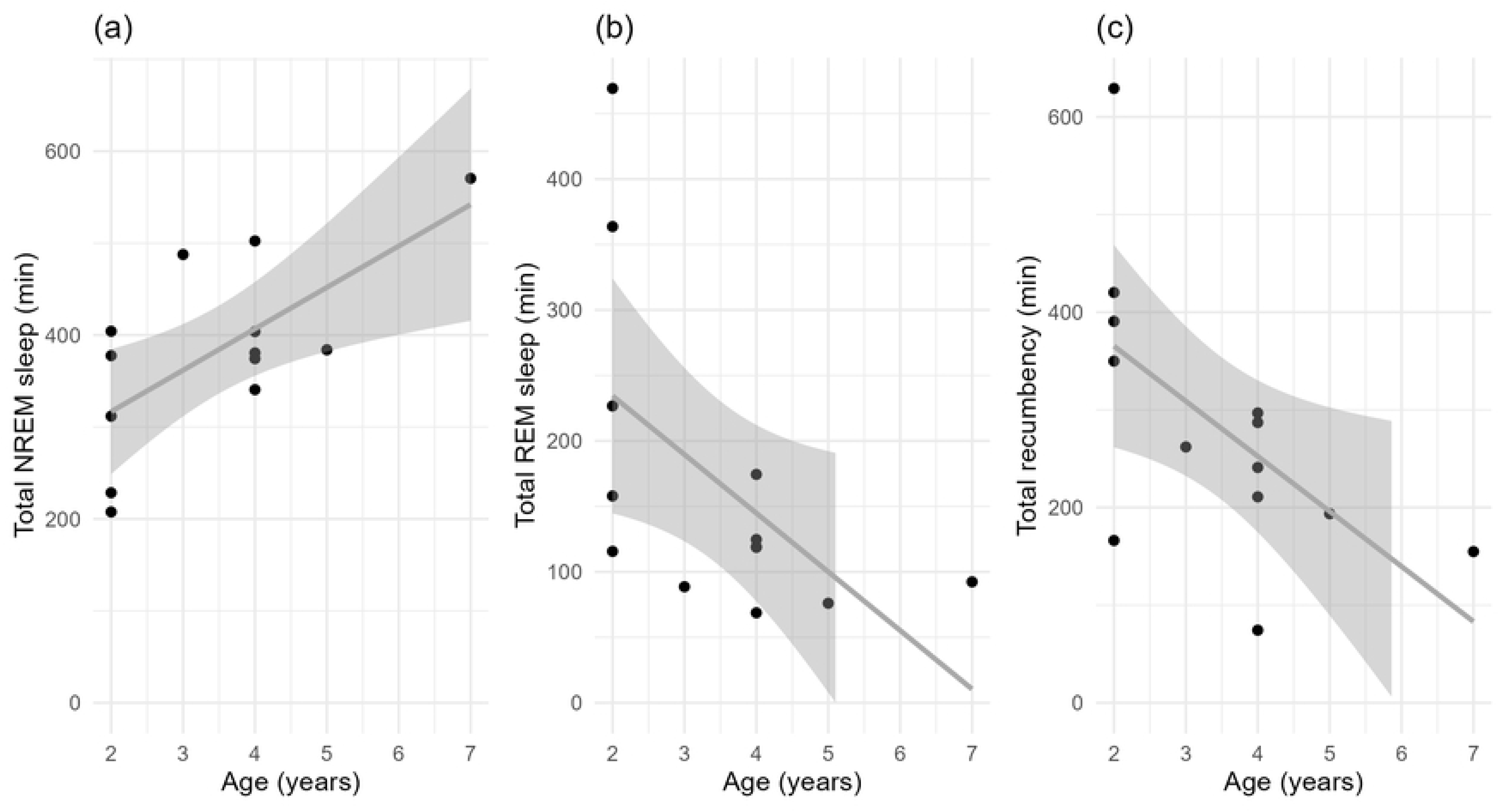
Associations between age and total NREM sleep and total REM sleep. A significant positive association was found with total NREM sleep (ANOVA: χ² = 5.4, p < 0.05), while a negative link was observed for total REM sleep (ANOVA: χ² = 4.4, p < 0.05) and total recumbency (ANOVA: χ² = 5.7, p < 0.05). Durations of sleep were obtained over a 48-hour period. The regression lines represent the model-predicted relationship between age and each sleep variable, while the shaded areas show the 95 % confidence intervals, indicating the precision of the estimates.

No significant associations were also found between TST and any of the three fixed factors (age, salivary cortisol and abnormal behaviours) in our study (Table 5).

#### Relationships between quality of sleep, age, salivary cortisol, and abnormal behaviours

Among the different SQIs, Total SQI and Combined SQI were significantly associated with at least one of the three studied factors (Table 5).

Specifically, Total SQI was negatively associated with the average salivary cortisol concentration (ANOVA: χ² = 5.3, p < 0.05; Fig. 3a) and the expression of abnormal behaviours (ANOVA: χ² = 4.1, p < 0.05; Fig. 4), whereas age was not significant. The marginal R² for this model was 37.9%. Similarly, Combined SQI was negatively associated with average cortisol concentration (ANOVA: F = 5.6, p < 0.05; Fig. 3b) but showed no significant relationship with abnormal behaviours or age. This model had an adjusted R² of 34.0%. Finally, neither Weighted Combined SQI nor NREM SQI and REM SQI showed any significant association with the three explanatory variables.

**Fig. 3:**
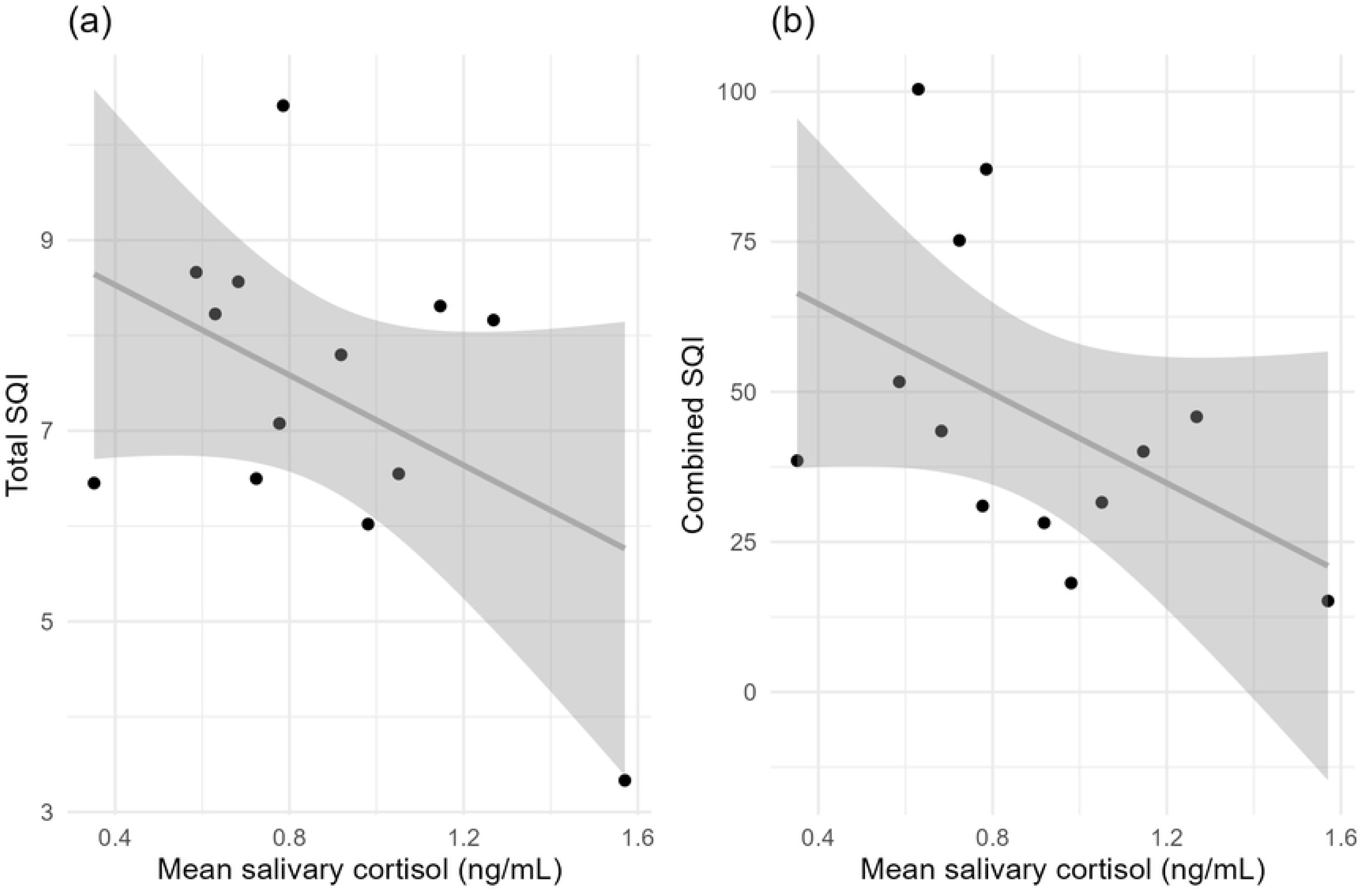
Associations between average salivary cortisol and a) duration of NREM sleep, b) Total SQI, c) Combined SQI and d) REM SQI. A significant negative association was found with Total and Combined SQIs (Total SQI: ANOVA: χ² = 5.3, p < 0.05; Combined SQI: ANOVA: F = 5.6, p < 0.05). Duration and qualities of sleep were calculated over a 48-hour period. The regression lines represent the model-predicted relationship between mean salivary cortisol and each sleep variable, while the shaded areas show the 95 % confidence intervals.

**Fig. 4:**
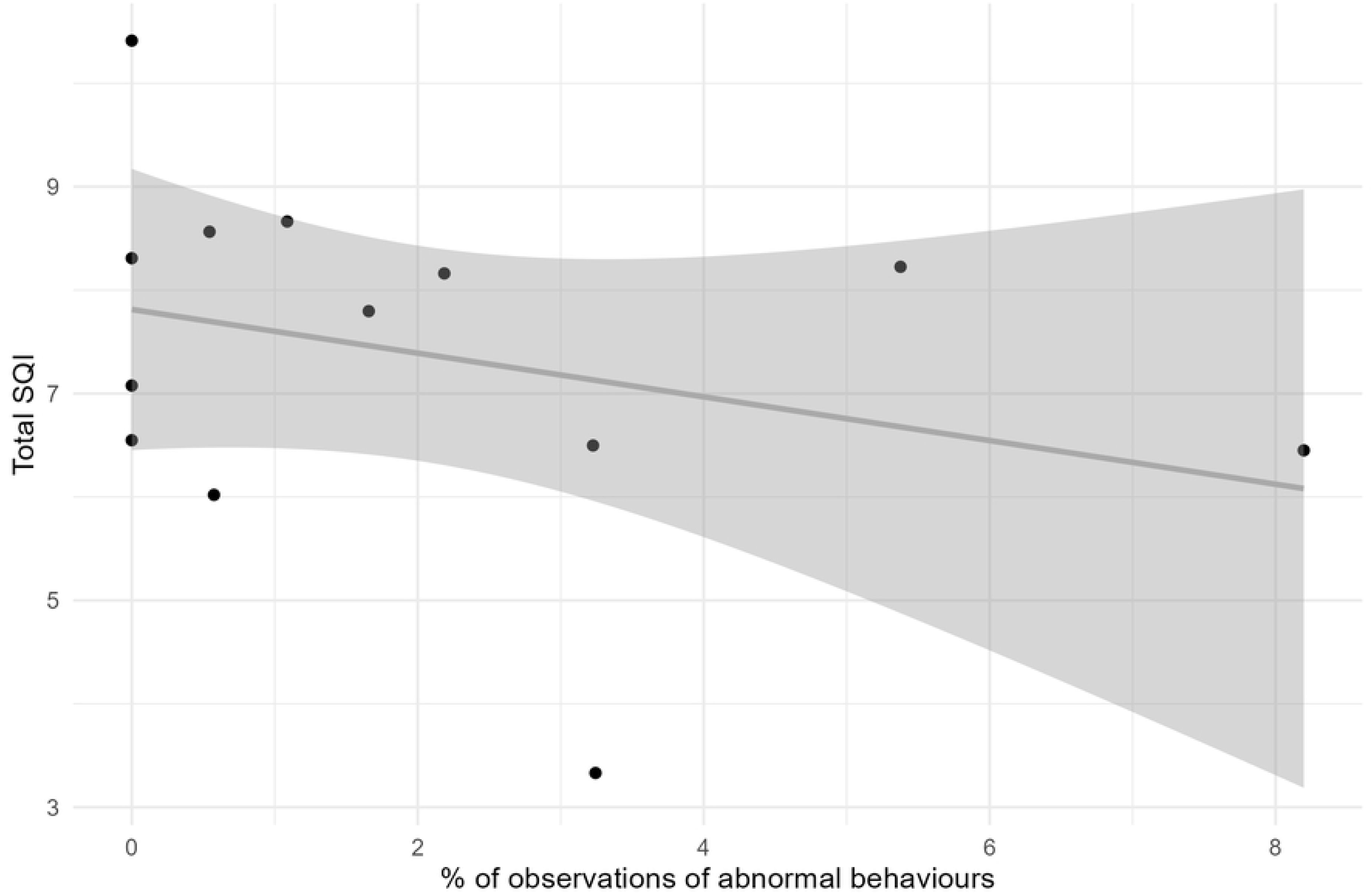
Association between percentage of observations of abnormal behaviours and Total SQI. A significant association was found (Anova: χ² = 4.1, p < 0.05). Total SQI and percentage of observations of abnormal behaviours of sleep were obtained over a 48-hour period. The regression line represents the model-predicted relationship between % of observations of abnormal behaviours and each sleep variable, while the shaded areas show the 95 % confidence intervals.

## Discussion

This field study represents the first investigation of sleep in racehorses, including both sleep quantity and quality, as well as its associations with age, morning cortisol levels, and abnormal behaviours. Results showed that sleep quantity (total NREM sleep, total REM sleep and total recumbency) was significantly associated with age, while sleep quality was linked to morning cortisol levels (Total SQI and Combined SQI) and the expression of abnormal behaviours (Total SQI). Overall, this study provides new insights into equine sleep patterns and highlights the link between sleep and welfare in racehorses.

### Descriptive overview of sleep, cortisol and abnormal behaviours in our population

#### Sleep

Horses in the present study slept on average 4.59 h/24 h, a value that is in the range of previous studies [1,20]. Indeed, it appears to be slightly higher than the 3.85 h/24 h reported in the review by Greening and McBride (2022), which compiled studies that measured sleep in horses using various methodologies. These slight differences between studies could be due to differences in environmental conditions, such as season or temperatures, bedding lighting regimes or to other individual characteristics of the horses, such as age, breed or physical activity.

REM sleep represented 30.6% of the TST in our study, which is higher than reported in other studies [1,20,34]. This difference may be explained by the fact that the horses in our sample were younger than those in the other studies. In many species, younger individuals that have not yet reached full maturity often require more REM sleep, partly because this sleep stage contributes to brain maturation [2]. Consistently, our own results showed that REM sleep decreased with age in the studied sample (see section 4.2).

Mean total lying time over 24 h was similar to that reported in a recent study on Lusitano horses [21], suggesting that posture-related rest patterns may be relatively consistent across certain breeds, despite differences in total sleep duration. However, a high degree of inter-individual variability was observed in our study, particularly for time spent in lateral recumbency. Because this is the most vulnerable sleeping posture, differences may reflect variation in stress sensitivity or perception of safety. These findings are consistent with the observations made by Griffin (2023), who found that individual differences in sensitivity to stress influenced recumbent sleep behaviours in horses with less recumbency in more sensitive individuals. However, in our study no relationship emerged between lying time and cortisol concentration or abnormal behaviours (see section 4.2.), suggesting that if personality or stress sensitivity underlies these differences, it was not captured by these indicators.

#### Salivary cortisol and abnormal behaviours

Salivary cortisol concentrations were within the expected range for horses at rest [36,37] and showed no day-to-day variation, supporting the use of mean values of cortisol for subsequent analyses exploring the relationship between morning cortisol levels and sleep patterns.

Abnormal behaviours were frequent with stereotypic behaviours observed in 7 out of 13 horses, representing 53.8% of the study population. While the prevalence of stereotypies is known to be high in Thoroughbred horses [27,38–40], the rate observed here was notably higher than that reported in other studies. This may be due to the characteristics of our sample or to the fact that behavioural scans were conducted across entire 24-h periods, including the night, which is rarely done and may capture stereotypic behaviours that are not expressed during the daytime. Additionally, the use of video recording eliminated potential observer effects on horses ‘behaviours. Alert and inactive behaviours were recorded at lower levels (respectively from 0 to 2.70%; and 0 to 3.83% of the total observations), but these were comparable to those reported in previous studies [27,41]. Finally, aggressive behaviours towards humans were rare (from 0 to 0.55% of the total observations), consistent with the literature, which found that Thoroughbred horses generally respond positively to human approach tests [27,40].

### Relationship between sleep, salivary cortisol, abnormal behaviours and age

Beyond these general patterns, we examined which factors could explain inter-individual variability in sleep, with a particular focus on age, morning cortisol as a marker of acute stress, and abnormal behaviours as indicators of chronic welfare impairment.

#### Quantity of sleep and age

Statistical analyses revealed that age significantly influenced sleep architecture. Specifically, total REM sleep decreased as expected and total NREM sleep increased with age, while TST remained stable. The significant negative association between age and total REM sleep, is consistent with observations in other mammalian species such as dogs [42] or humans [3], where REM sleep tends to decline with age, potentially due to age-related alterations in the cholinergic system that play a key role in REM initiation and maintenance [43]. Interestingly, this redistribution of sleep architecture suggests that, while older horses do not sleep more overall, they may shift towards increased NREM sleep, possibly reflecting a greater physiological need for recovery, at the expense of REM functions. Remarkably, this trend was already detectable within the relatively narrow age range of our sample (2–7 years; mean: 3.3 years).

Additionally, a significant positive association was found between age and total NREM sleep duration. In humans, both TST and NREM sleep tend to decrease with age, however, some species, such as mice, show the opposite pattern, with TST and NREM sleep increasing with age, particularly during their active phase [44]. These interspecies differences raise the question of whether age-related sleep changes are species-dependent [45]. In our sample, horses appeared to follow a pattern more similar to those reported in rodents than humans, with older individuals exhibiting longer NREM sleep duration. This interpretation aligns with the observation that NREM sleep proportion was lower in our younger sample compared to Greening et al. (2021), where horses were older. Moreover, because horses in the current study were athletes engaged in regular training and racing, they were likely to accumulate greater physical strain and potential muscle microtrauma with increasing age [46]. The observed increase in NREM sleep with age could reflect a greater physiological need for recovery. This hypothesis is supported by the role of NREM sleep in promoting growth hormone secretion, which contributes to physical restoration [47,48]. As growth hormone release declines with age in humans [49] and horses [50], older horses might require longer NREM periods to achieve equivalent recovery. Since TST did not differ with age in our study, this pattern could suggest a redistribution of sleep architecture, favouring NREM at the expense of REM.

The increase in NREM sleep and the decrease in REM sleep with age are consistent with our finding that age was also negatively associated with total recumbency duration. Because recumbency (as a result of atonia) is essential for achieving REM sleep in horses [22], shorter recumbency duration may limit the opportunity to enter or maintain REM sleep. Conversely, individuals that spend more time in recumbency may have an increased probability of reaching REM sleep, thus sustaining longer REM periods.

#### Quality of sleep, salivary cortisol and abnormal behaviours

Our findings indicate that both acute stress (morning cortisol) and chronic welfare indicators (abnormal behaviours) were associated with sleep quality. Specifically, higher cortisol concentrations were linked to lower total and combined SQI, while a greater expression of abnormal behaviours was only associated with lower total SQI. The association between elevated cortisol and sleep fragmentation aligns with previous findings in other species: for example, in young children, more fragmented sleep was associated with higher morning cortisol concentrations [51], and in adults with a lower early decline of salivary cortisol [52].

Moreover, the positive relationship between abnormal behaviours and total SQI suggests that there is a link between poor quality sleep and poor welfare [26,53]. In both humans and animals, stress, anxiety, and apathy have been linked to sleep disturbances [54–56]. However, it is also known that poor sleep can also enhance levels of stress and reduce welfare [57,58]. Thus, it is important to note that a correlation of poor sleep with markers of stress and poor welfare does not imply causality in a specific direction, but highlights a reciprocal relationship whereby stress, welfare and sleep disturbances are intertwined.

### Study limitations

These results stem from an initial field investigation, in which not all parameters could be fully controlled. Other factors, such as sex, could also influence sleep patterns, but the limited sample size, reflecting practical constraints since manual sleep analysis is time-consuming, did not allow their inclusion in our statistical models.

## Conclusions

This study represents the first detailed description of the sleep of training Thoroughbred horses, exploring both the quality and quantity of sleep and their relationships with age, abnormal behaviours (i.e., alertness, stereotypy, inactivity, and aggressiveness towards humans), and cortisol. Quantity of sleep was associated with age, and specifically positively associated with total NREM sleep and negatively associated with total REM sleep, suggesting that sleep patterns in the horse, like other mammalian species, evolve with age, favouring NREM over REM in older individuals. Additionally, sleep quality seems to be negatively related with morning salivary cortisol concentration (Total and Combined SQI) and the expression of abnormal behaviours (Total SQI). Together, these relationships suggest a possible link between higher stress levels or welfare impairment and poorer sleep quality.

These findings highlight the importance of management strategies that not only ensure adequate housing conditions but also minimise stress and support individual recovery capacities. While all horses in this study were kept under calm conditions with deep bedding, known to improve good sleep (Greening et al., 2021, 2013; Pedersen et al., 2004), differences in sleep quality still emerged, suggesting that other factors may play a crucial role in promoting restorative sleep. These factors might include training load, recovery periods, opportunities for social contact, and environmental predictability.

As poor sleep has been linked to fatigue, mood alterations, injury risk and performance deficits in human athletes [5,62], ensuring high-quality sleep in racehorses may help to reduce stress, prevent abnormal behaviours from developing or becoming worse, and ultimately improve overall welfare and performances.

## Acknowledgements

The authors would like to thank the trainers, riders, and stable staff for their cooperation and support during data collection. We are also grateful to Camille Mikaeloff for her assistance with the study and for performing the blood sampling.

## Notes

### Competing Interest Statement

The authors have declared no competing interest.

